# A quantitative modular modeling approach reveals the consequences of different A20 feedback implementations for the NF-kB signaling dynamics

**DOI:** 10.1101/582767

**Authors:** Janina Mothes, Inbal Ipenberg, Seda Çöl Arslan, Uwe Benary, Claus Scheidereit, Jana Wolf

**Affiliations:** Mathematical Modelling of Cellular Processes, Max Delbrück Center for Molecular Medicine, Berlin, Germany; Signal Transduction in Tumor Cells, Max Delbrück Center for Molecular Medicine, Berlin, Germany

**Author notes:** Institut für Neuroimmunologie und Multiple Sklerose, Zentrum für Molekulare Neurobiologie Hamburg, Universitätsklinikum Hamburg-Eppendorf, Hamburg, Germany (Inbal Ipenberg). Corresponding author (JW).

**Keywords:** quantitative modelling, modular modelling, feedback, regulation, NF-κB signalling, A20, IKK regulation, response time, signalling dynamics

## Abstract

Signaling pathways involve complex molecular interactions and are controlled by non-linear regulatory mechanisms. If details of regulatory mechanisms are not fully elucidated, they can be implemented by different, equally reasonable mathematical representations in computational models. The study presented here focusses on NF-κB signaling, which is regulated by negative feedbacks via IκBα and A20. A20 inhibits NF-κB activation indirectly through interference with proteins that transduce the signal from the TNF receptor complex to activate the IκB kinase (IKK) complex. We focus on the question how different implementations of the A20 feedback impact the dynamics of NF-κB. To this end, we develop a modular modeling approach that allows combining previously published A20 modules with a common pathway core module. The resulting models are based on a comprehensive experimental data set and therefore show quantitatively comparable NF-κB dynamics. Based on defined measures for the initial and long-term behavior we analyze the effects of a wide range of changes in the A20 feedback strength, the IκBα feedback strength and the TNFα stimulation strength on NF-κB dynamics. This shows similarities between the models but also model-specific differences. In particular, the A20 feedback strength and the TNFα stimulation strength affect initial and long-term NF-κB concentrations differently in the analyzed models. We validated our model predictions experimentally by varying TNFα concentrations applied to HeLa cells. These time course data indicate that only one of the A20 feedback models appropriately describes the impact of A20 on the NF-κB dynamics.

**Author summary:** Models are abstractions of reality and simplify a complex biological process to its essential components and regulations while preserving its particular spatial-temporal characteristics. Modelling of biological processes is based on assumptions, in part to implement the necessary simplifications but also to cope with missing knowledge and experimental information. In consequence, biological processes have been implemented by different, equally reasonable mathematical representations in computational models. We here focus on the NF-κB signaling pathway and develop a modular modeling approach to investigate how different implementations of a negative feedback regulation impact the dynamical behavior of a computational model. Our analysis shows similarities of the models with different implementations but also reveals implementation-specific differences. The identified differences are used to design and perform informative experiments that elucidate unknown details of the regulatory feedback mechanism.

## Introduction

Transcription factor NF-κB regulates cell differentiation, proliferation and survival. In line with its broad range of normal physiological functions, aberrant activation of NF-κB can lead to severe diseases, e.g. autoimmune, neurodegenerative and cardiovascular diseases as well as cancer and diabetes (1, 2). In resting cells, the transcription factor NF-κB is located in the cytoplasm bound to IκBα, which prevents the translocation of NF-κB into the nucleus. Upon stimulation, e.g. with TNFα, the IκB kinase (IKK) complex is activated. The IKK complex phosphorylates IκBα, marking it for proteasomal degradation. Released NF-κB translocates into the nucleus and activates the transcription of a number of target genes (3). Two of these are NFKBIA, encoding IκBα, and TNFAIP3, encoding A20. Both proteins exhibit negative feedbacks on NF-κB activation. IκBα binds to NF-κB retrieving it from the DNA and thus exhibiting a direct negative feedback (4). A20 inhibits NF-κB activity indirectly through interference with proteins mediating the signal from the TNF receptor complex to the IKK complex (5). The exact molecular mechanism of the inhibitory effect of A20 on the IKK complex is still under discussion (6-8).

In the last decades, several mathematical models describing the NF-κB signaling have been published (9-15), and reviewed (16-19). All models comprise the core processes of the canonical NF-κB signaling, e.g. the interaction of NF-κB and IκBα and the transcription and translation of IκBα as well as the IKK-induced degradation of IκBα. The majority of those models include only the negative feedback via IκBα, which has been well-studied and characterized (14).

Until today, only a small number of mathematical models has been developed that include the A20-dependent negative feedback mechanism (10, 13, 20, 21). These models utilize similar implementations of the core signaling processes but differ in their implementation of the A20 feedback. Since the exact inhibitory mechanism of A20 on IKK has not yet been fully elucidated, the models implement different hypotheses. While the model of Lipniacki et al. (2004) (10) and the derived model by Ashall et al. (2009) (21) implement the inhibitory action of A20 on the level of IKK, the models of Werner et al. (2008) (20) and Murakawa et al. (2015) (13) basically implement the hypothesis that A20 blocks the signaling upstream of IKK by binding to TNF receptor associated proteins. In particular, the models by Lipniacki et al. (2004) and Ashall et al. (2009) comprise three different states of IKK: neutral, active and inactive. In the model proposed by Lipniacki et al. (2004), A20 promotes the inactivation of activated IKK, whereas, in the model by Ashall et al. (2009) A20 inhibits the ‘recycling’ of inactive IKK to neutral IKK and consequently the activation of IKK. In the models by Werner et al. (2008) and Murakawa et al. (2015), A20 inhibits basal and TNFα-induced IKK activation, although Werner et al. (2008) consider the signaling mechanisms upstream of IKK with substantially more molecular detail than Murakawa et al. (2015). In short, all four models share a feedback inhibition of IKK activity by A20 but differ in the specifics of their A20 feedback implementations.

Here, we compare the different A20 feedback structures. We selected those implemented in the models of Lipniacki et al. (2004) (10), Ashall et al. (2009) (21), and Murakawa et al. (2015) (13), because these capture three different hypotheses and the models are comparable at their level of detailedness. We did not include the model of Werner et al. (2008) because its A20 feedback mechanism is essentially captured with reduced complexity in the model of Murakawa et al. (2015). We addressed the question whether the different feedback implementations affect NF-κB dynamics in similar or distinct ways. To this end, we used a computational approach in which we established three ordinary differential equation (ODE) models. Each model is composed of a core module and an upstream module (Fig 1A). The core module is identical in all three models and describes the interaction of NF-κB and IκBα, transcription and translation of IκBα, and IKK-induced degradation of IκBα. The three upstream modules comprise the three distinct mechanisms of IKK inhibition by A20 that Lipniacki et al. (2004), Ashall et al. (2009) and Murakawa et al. (2015) have proposed. In this way, we used a modular concept to derive three models that share an identical core module but differ in their implementations of the A20 feedback in the upstream module. By fitting these models to the same set of experimental data, we derive models showing quantitatively similar NF-κB dynamics. We use this approach to directly compare the influences of the structural difference in the upstream modules on the response of the NF-κB dynamics. In particular, we focused on the impact of the A20 and IκBα feedback strength. Moreover, we analyze in each model how the A20 feedback modulates the effect of varied TNFα stimulations on the NF-κB dynamics. We find that the different A20 feedback implementations exert similar but also model-specific effects and use the predicted distinct dynamic responses towards incremental alterations of TNFα stimulation strength for an experimental validation of our results.

**Fig 1:**
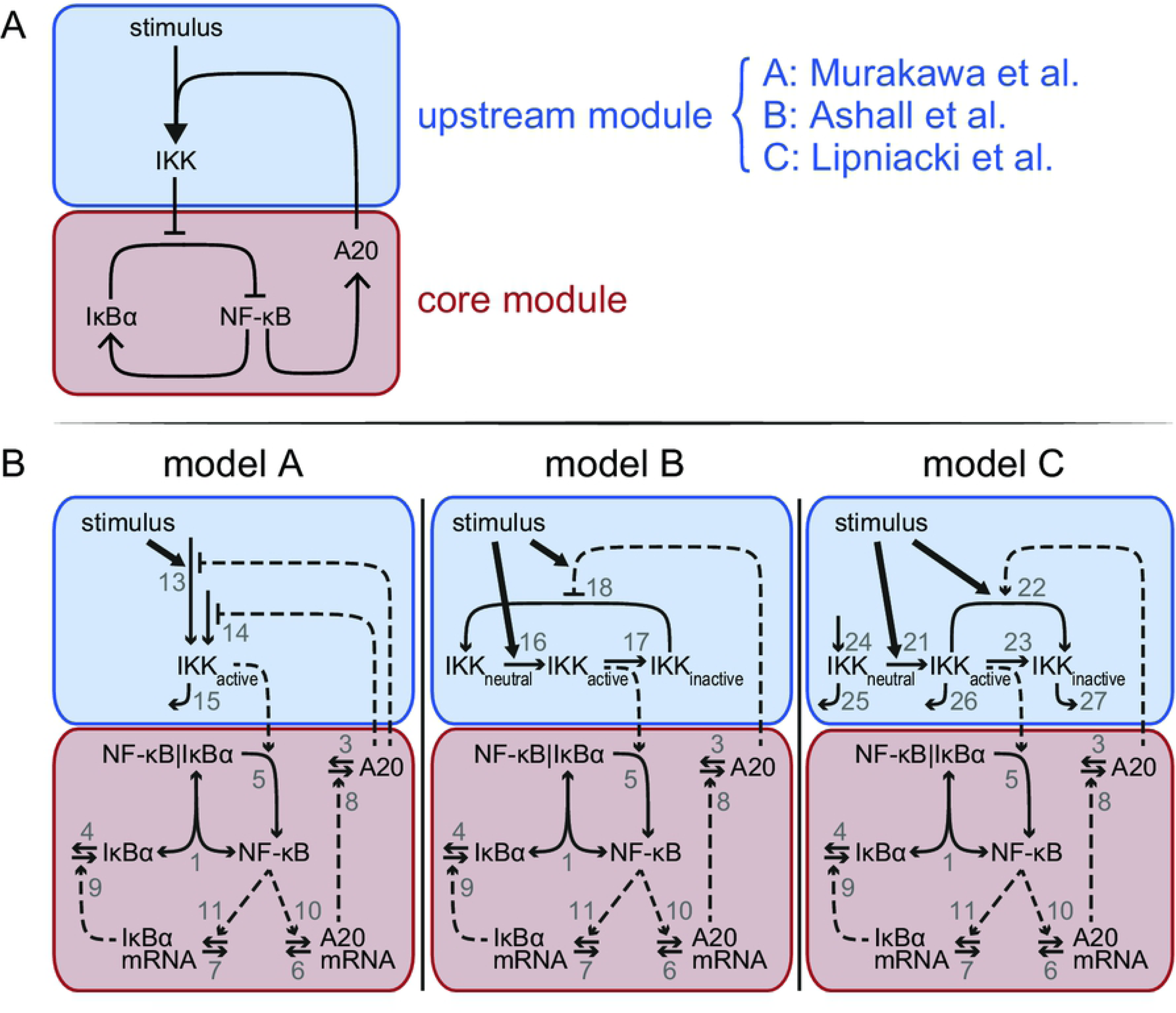
Model schemes comprising the common core module and distinct upstream modules. A: Each model is composed of a core module (red) and an upstream module (blue). The core module is identical in each model but the upstream module differs between model A, B, and C, implementing the A20 feedback mechanisms proposed by (13), (21) and (10), respectively. B: Schematic representations of the three models A-C. Vertical bars separate components in a complex. One-headed arrows indicate the direction of the reaction; double-headed arrows illustrate reversible binding reactions. Dashed arrows represent activation processes; the dashed lines ending in T-shape denote inhibition. The number next to an arrow specifies the number of the reaction. Model equations and the reference parameters are provided in S1 File.

## Methods

### Model structures

In order to compare the three distinct implementations of the inhibitory mechanism of A20, we modularly designed three models. These models comprise an identical core module to which different upstream modules are attached (Fig 1A, B). The upstream modules are those proposed by Lipniacki et al. (2004) (10), Ashall et al. (2009) (21) and Murakawa et al. (2015) (13) capturing the different A20 feedback implementations. The overall models are hereafter referred to as model A, B and C

The common core module of models A-C (Fig 1B) describes the reversible binding of free NF-κB and IκBα (reaction 1). Activated IKK (IKK_active_) induces the IκBα degradation releasing NF-κB from the complex (reaction 5). Unbound NF-κB induces the transcription of IκBα mRNA (reaction 11), which is translated to IκBα (reactions 9). IκBα mRNA and IκBα protein degrade via reactions 7 and 4, respectively. In addition to IκBα mRNA, NF-κB induces the transcription of A20 mRNA (reaction 10). A20 mRNA is translated to A20 (reaction 8). A20 mRNA and protein are degraded via reactions 6 and 3, respectively. Taken together, the core module consists of five ordinary differential equations (ODEs) and one conservation relation for NF-κB. A detailed description of the corresponding rates and a list of the parameters are provided in S1 File.

The upstream module of model A (Fig 1B, left) comprises a very condensed representation of the activation of the IKK complex. The abundance of IKK_active_ increases in a TNFα-dependent and independent manner (reactions 13 and 14, respectively), both of which are inhibited by A20. IKK_active_ is inactivated via reaction 15. In the upstream module of model B (Fig 1B, middle), IKK cycles between three distinct states: IKK_neutral_, IKK_active_, and IKK_inactive_. TNFα stimulation converts IKK_neutral_ into IKK_active_ (reaction 16), IKK_active_ is converted to IKK_inactive_ (reaction 17) and IKK_inactive_ is finally turned over to IKK_neutral_ again (reaction 18). A20 inhibits this last reaction in a stimulus-sensitive manner.

The upstream module of model C (Fig 1B, right) includes the same states of IKK as described in model B, but IKK_neutral_, IKK_active_, and IKK_inactive_ do not interconvert in a cycle, i.e. obey a conservation relation. Instead, IKK_neutral_ is continuously produced (reaction 24) and all three forms of IKK are subject to degradation (reactions 25-27). Similar to model B, TNFα stimulation in model C also converts IKK_neutral_ into IKK_active_ (reaction 21), which in turn forms IKK_inactive_ (reaction 23). In contrast to model B, model C includes an additional mechanism to convert IKK_active_ into IKK_inactive_ (reaction 22). TNFα stimulation as well as A20 enhance this conversion.

Taken together, model A consist of one ODE in its upstream module in addition to the five ODEs and one conservation relation of NF-κB in the core module; model B incorporates two additional ODEs and an additional conservation relation of IKK in the upstream module; and model C includes three additional ODEs in its upstream module. Detailed descriptions of all three models are given in S1 File.

### Model parametrizations

To parameterize the ODEs of the core module, we decided to use the parameters from our previously published model (13). This approach was based on two arguments. First, this model is based on a comprehensive data set characterizing the modulation of A20 feedback strength and its impact on NF-κB dynamics. Secondly, the core processes of this model perfectly match the reactions of the core module of our models A-C.

To parameterize the three different upstream modules of models A-C, we initially used the parameters published for the corresponding models (10, 13, 21). However, simulations of models A-C showed very diverse dynamics of unbound NF-κB in response to identical TNFα stimulation conditions (Fig 2A). For instance, the concentration of free NF-κB transiently increases in models A and B, but on a slower time scale in model A. In contrast, unbound NF-κB hardly increases upon TNFα stimulation in model C.

**Fig 2:**
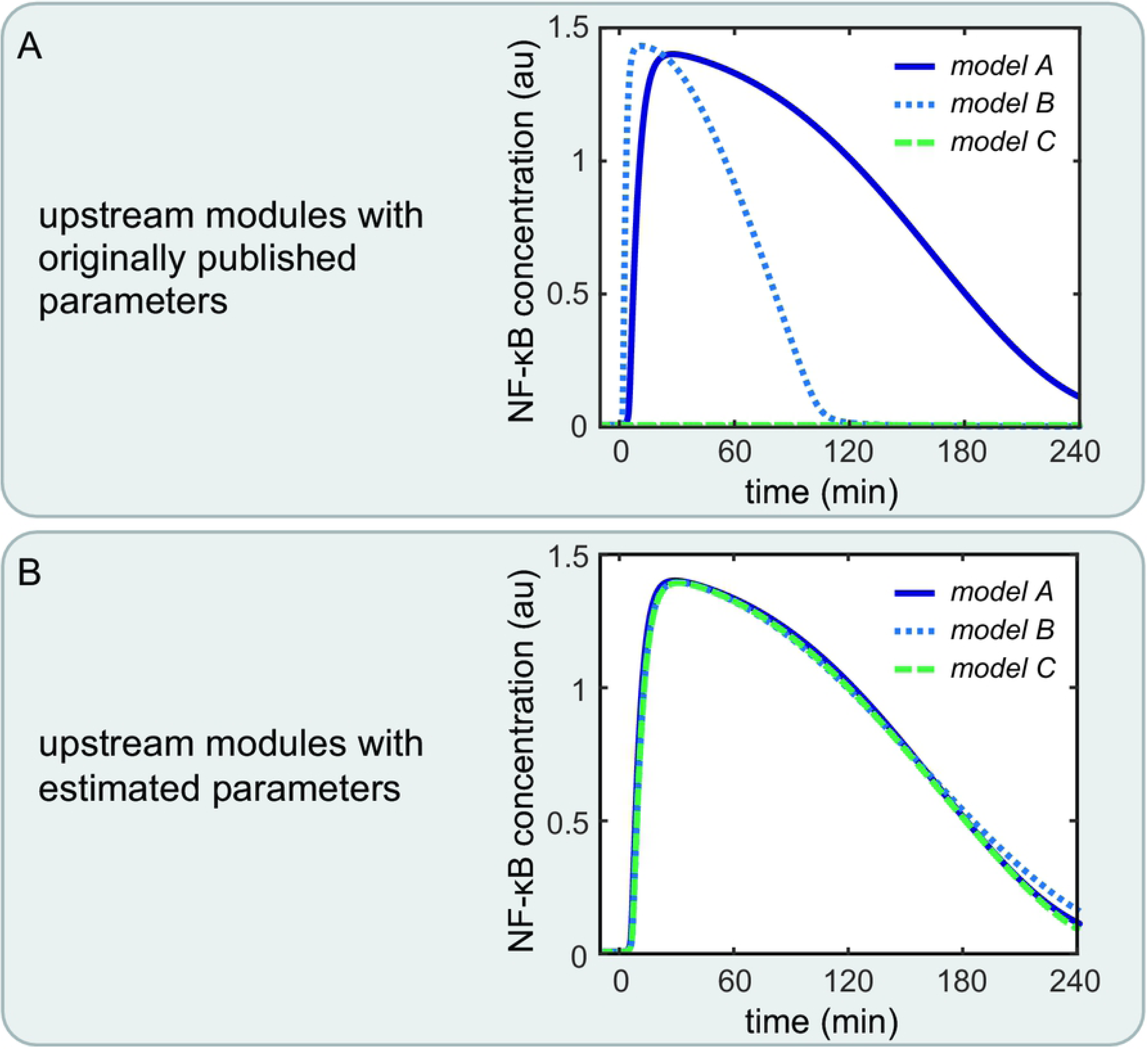
NF-κB dynamics of the three models comprising the core module and the indicated upstream module. A: Differences in NF-κB dynamics can be observed for the three models using the originally published parameters. B: Nearly identical NF-κB dynamics can be observed for the three models with newly estimated parameters for the upstream modules.

In order to compare models A-C directly, it is necessary that NF-κB exhibits the same dynamics upon TNFα stimulation in all three models. Thus, we estimated new parameters of the reactions in the upstream modules such that all components of the core module show the same dynamics in all three models. We used the D2D Toolbox (22) to estimate these parameters while keeping the parameters of the core module fixed. With this restriction on the parameters of the core module, we were able to reasonably minimize the parameter search space and obtain identical dynamics of the components of the core module. The details of the parameter estimation are explained in S1 File. Simulations of models A-C with these estimated parameters showed nearly identical dynamics of NF-κB activation upon TNFα stimulation (Fig 2B) and all remaining components of the core module (Figs 1 and 2 in S1 File).

Next, we checked whether the new parameterization changed the inhibitory effect of A20 on the activation of IKK. To do so, we simulated A20 knockout conditions by setting the A20 transcription rate k_10_ to zero and compared the resulting dynamics to those of wild-type conditions, i.e. using the reference value of k_10_ (Table 1 in S1 File). The simulations show that the A20 knockout causes a prolonged increase in NF-κB, IKK and IκBα mRNA upon TNFα stimulation compared to wild-type (23) in all three models (Figs 3-5 in S1 File). The simulations furthermore show that the absence of A20 leads to a decrease in IκBα concentration in all three models. These results demonstrate that the parameterizations of the models A-C do represent the inhibitory effect of A20 on the activation of IKK.

Taken together, models A, B and C were derived by modular design from an identical core module and different upstream modules specifying distinct implementations of the A20 feedback and TNFα stimulation. The models exhibit almost identical dynamics of their common model components, and show similar dynamical behavior in A20 knockout simulations.

### Quantitative characterization of the NF-κB dynamics

To quantitatively compare the dynamics of unbound NF-κB between the models A-C, we characterized NF-κB dynamics by three measures (Fig 3): (1) the maximal NF-κB concentration (x_max_), (2) the time of the maximal NF-κB concentration (t_max_), and (3) the response time (t_r_,) defined in (24), which quantifies the time required for a complete NF-κB response after stimulation. While x_max_ and t_max_ describe the initial response of NF-κB to TNFα stimulation, t_r_ represents a normalized duration of NF-κB signaling and can therefore be used as a measure for the long-term dynamics.

**Fig 3:**
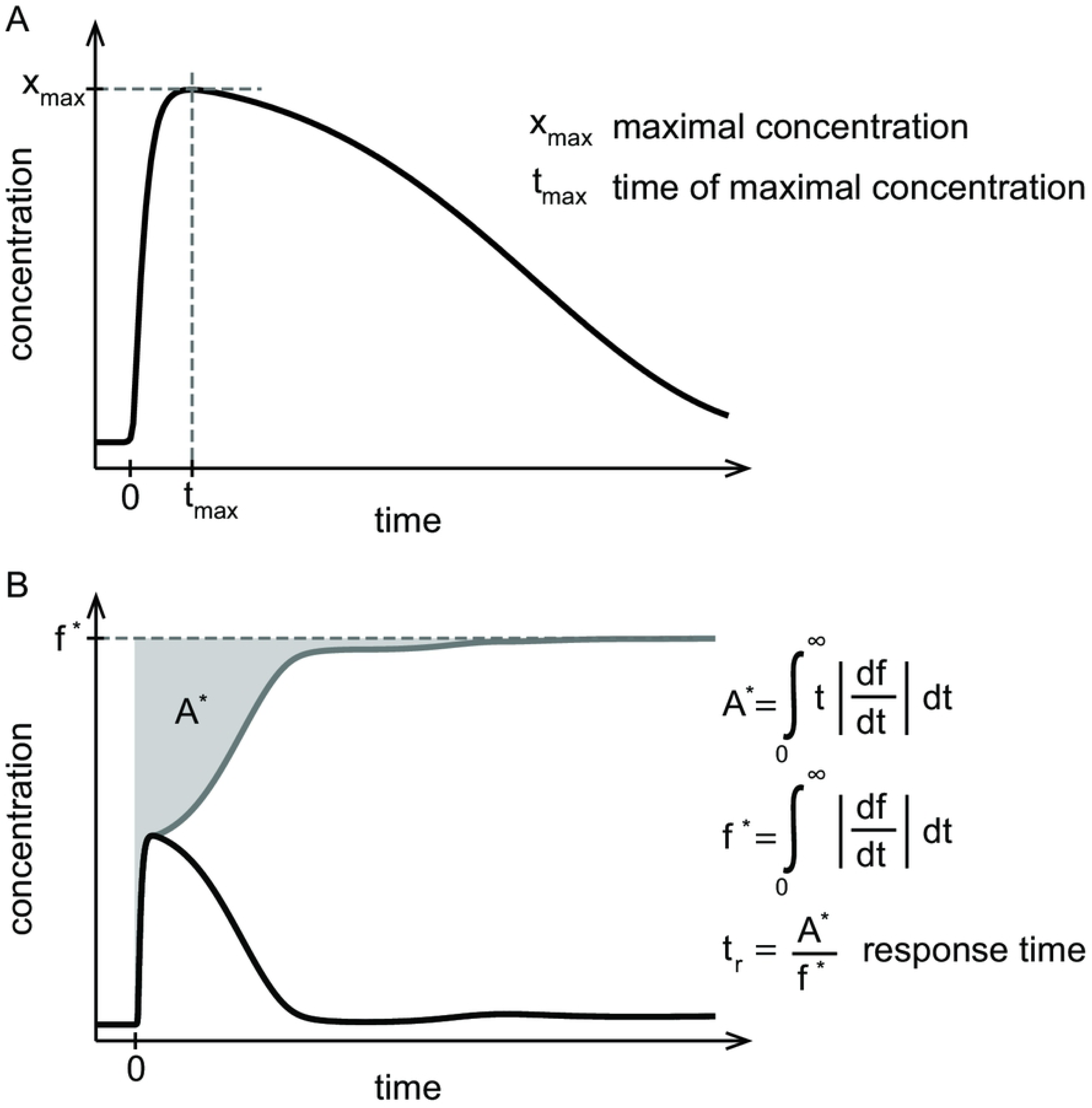
Measures to quantify NF-κB dynamics. A: The maximal concentration of NF-κB (x_max_) and the time of the maximal concentration of NF-κB (t_max_) characterize the initial NF-κB response. B: The response time (t_r_) defined in (24) is determined by the grey area (A*) normalized to the steady state (f*) of the absolute gradient of the dynamics of NF-κB. The response time quantifies the time required for the activation and deactivation of NF-κB upon stimulation and can be interpreted as a characterization of the NF-κB long-term behavior.

### Numerical simulations

The model equations are listed in S1 File. Calculations were done with MathWorks Matlab R2013b. Steady state solutions were numerically obtained. Starting from those steady state solutions, the models are always simulated for 57600 min.

### Experimental methods

HeLa cells were stimulated with 10, 25 or 100 ng/ml TNFα (human recombinant TNFα, Alexis Corporation) for the time periods indicated (120, 100, 80, 60, 40, 20, 10 min) or were left untreated. Following stimulation, cells were lysed in 20 mM Hepes pH=7.9, 450 mM NaCl, 1 mM MgCl2, 0.5 mM EDTA pH=8.0, 0.1 mM EGTA, 1% NP-40, 20% glycerol, supplemented with complete protease inhibitor mixture and Phosphostop (Roche Applied Science), 50 nM Calyculin A, 10 mM NaF, 10 mM β-glycerophosphate, 0.3 mM Na3VO4 and 1 mM Dithiothreitol. Lysates were centrifuged at 14,000 rpm for 10 min.

NF-κB DNA-binding activity was assayed by Electrophoretic Mobility Shift Assay (EMSA) as previously described (25).

EMSA quantification was made using the phosphor-imager Typhoon FLA 9500, GE Healthcare. Data were quantified using ImageQuant software. After background subtraction, the NF-κB band was normalized to a respective constant non-specific band.

## Results

### Effects of different A20 feedback strengths on NF-κB dynamics

As a starting point, we studied the impact of the A20 feedback on the NF-κB dynamics upon a constant TNFα stimulation. To do so, we varied the A20 feedback strength and studied its effects on the temporal change of the concentration of unbound NF-κB (hereafter denoted NF-κB) in all models. The strength of the A20 feedback is varied by multiplying the transcription rate constant of the A20 mRNA (k10) with a factor, i.e. feedback strength. A low value of the feedback strength corresponds to a weak negative feedback, whereas a high feedback strength results in a strong negative feedback. Local sensitivity analyses showed that a variation of the translation rate constants of A20 (k8) and of the transcription rate constant have a comparable effect on the three measures of the NF-κB dynamics (Figs 6-8 in S1 File). Thus, our choice to vary the transcription rate constant by a factor, i.e. the feedback strength, rather than the translation rate constant does not affect our conclusions.

The NF-κB dynamics of the models A-C for the A20 feedback strengths 0.1 and 10 are shown in Fig 4A. In case of a high A20 feedback strength of factor 10, models B and C show a fast and transient increase of NF-κB concentration upon a constant TNFα stimulation (Fig 4A – top). In model A, NF-κB increases later and to a lesser extent compared to model B and C, yet it decreases to a similar final concentration. In the case of a low A20 feedback strength of factor 0.1 (Fig 4A – bottom), all three models show an almost identical increase in the NF-κB concentration. However, NF-κB decreases faster and to a lower final concentration in model C compared to model A and B. Comparing the simulations of the high with the low A20 feedback strength, all three models show a faster decrease in NF-κB in the case of high compared with low A20 feedback strength.

**Fig 4:**
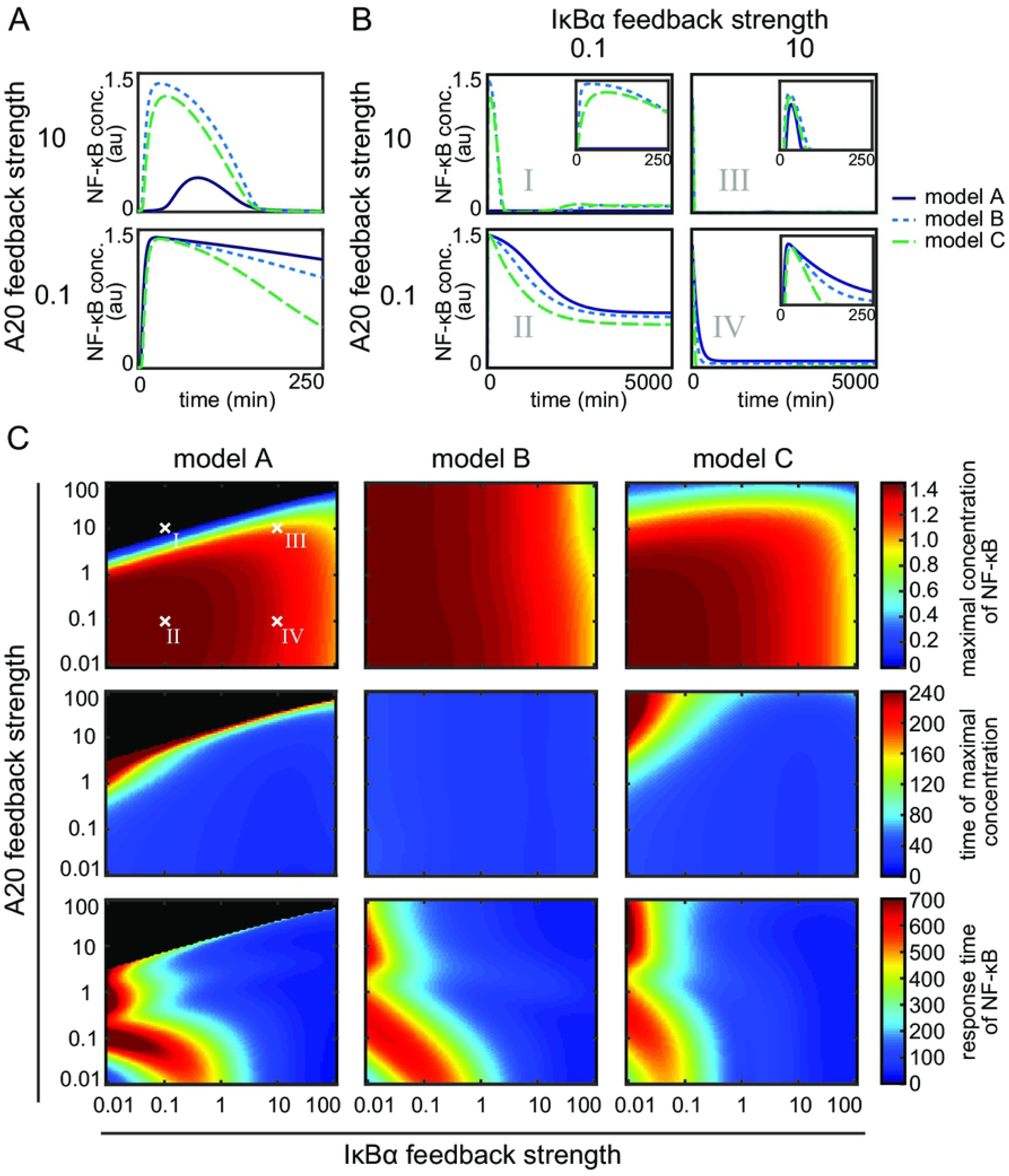
Influence of the A20 feedback strength and the IκBα feedback strength on NF-κB dynamics. A: NF-κB dynamics of the three models for two different A20 feedback strengths. B: NF-κB dynamics of the three models for four exemplary combinations of A20 and IκBα feedback strengths. C: The effect of the different combinations of feedback strengths on the maximal concentration of NF-κB (first row), the time of the maximal concentration (second row) and the response time of NF-κB (third row) in the case of model A (first column), model B (second column) and model C (third column). The four exemplary combinations of feedback strength shown in panel B (I, II, III, and IV) are indicated. Black areas mark the combinations of feedback strengths where hardly any NF-κB response is observed, i.e. the difference between maximal concentration of NF-κB and initial concentration of NF-κB is less than the threshold value of 0.001 μM.

These results reflect the strong influence of the A20 feedback on the deactivation of NF-κB. A high A20 feedback strength causes a stronger and faster deactivation in all three models. Moreover, in model A a strong A20 feedback strength notably reduces and also delays NF-κB activation.

### The IκBα feedback modulates the effect of the A20 feedback on NF-κB

Besides A20, IκBα is an important negative regulator of NF-κB dynamics. We next analyzed whether the interplay of these two feedbacks in the regulation of NF-κB dynamics is similar in the three models. To address this question, we varied the IκBα feedback strength in addition to that of A20. Similar to the A20 feedback strength, we multiplied the transcription rate constant of the IκBα mRNA (k11) by a factor to change the IκBα feedback strength.

The NF-κB dynamics of the three models for four exemplary combinations of different A20 and IκBα feedback strengths are shown in Fig 4B (cases I – IV). The simulations show a rapid increase of NF-κB concentration upon TNFα stimulation for all models and in all four cases (I-IV), with one exception (model A, case I). The subsequent decrease of NF-κB concentration differs in strength and pace. For a combination of a high A20 feedback strength and a low IκBα feedback strength (case I), NF-κB concentrations in models B and C decrease to the half-maximum level at around 250 min whereas model A shows no NF-κB response to TNFα stimulation. When A20 and IκBα feedback strength are both low (case II), NF-κB concentration decreases at a much slower pace and to lesser extent than in case I for models B and C; here (case II) model A also shows a transient NF-κB activation. If the feedback strengths of A20 and IκBα are high (case III), a fast increase can be observed that is followed by a nearly complete decrease of NF-κB concentration at 100 min for all models. For combinations of a high IκBα feedback strength with a low A20 feedback strength (case IV), the decrease in NF-κB concentration is slightly prolonged compared to case III, depending also on the model. These results are in agreement with our earlier finding that higher A20 feedback strengths cause a faster and stronger decrease in NF-κB than lower A20 feedback strengths (Fig 4A).

In the comparison of case I and case III, which both comprise the same A20 feedback strength but differ in their IκBα feedback strength, a stronger as well as faster decrease in the NF-κB concentration can be observed for high IκBα feedback strengths. The comparison of case II and case IV yields a similar result, showing that a higher IκBα feedback strength leads to a faster and stronger decrease in NF-κB concentrations and therefore influencing its short-term and long-term dynamics.

In summary, both feedbacks lead to the deactivation of NF-κB after a transient increase. Thus, if only one of the two feedbacks is strong, it can compensate for the other. If A20 and IκBα feedback strengths are both strong, the effect on the deactivation of NF-κB is enhanced resulting in an even faster and stronger NF-κB deactivation.

Beside these general observations, we find model-specific effects of the feedbacks. Most obviously, the maximal NF-κB activation and the deactivation pace seem to vary between the models. An interesting combination is a strong A20 with a low IκBα feedback strength (case I) for model A, which prevents an NF-κB response to TNFα stimulation.

### Quantification of the influences of the A20 and the IκBα feedback on NF-κB dynamics

To determine to what extent the models A-C differ in their NF-κB response under the various feedback strengths, we quantified the dynamics of NF-κB by three measures: the maximal concentration of NF-κB, the time of the maximal concentration, and the response time (Fig 3). The first two measures characterize the initial NF-κB dynamics whereas the last measure characterizes the long-term NF-κB dynamics. For each model we then continuously varied the A20 and the IκBα feedback strengths over a broad range of four orders of magnitude, covering very low (e.g. 0.01) as well as very high (e.g. 100) feedback strengths (Fig 4C).

In model A, the maximal NF-κB concentration barely changes at A20 feedback strengths below 1 (Fig 4C – first column, first row). In those cases, only an increase in the IκBα feedback strength leads to a decrease in the maximal concentration of NF-κB. For strong A20 feedback strengths above 1, the A20 feedback can prevent the NF-κB response almost completely for a wide range of different IκBα feedback strengths (Fig 4C – first row, black area). This is in agreement with case I in Fig 4B showing no NF-κB response for high A20 and low IκBα feedback strengths. For A20 feedback strengths below 1 in combination with a wide range of different IκBα feedback strengths, the maximal concentration of NF-κB is reached in the first 80 min (Fig 4C – first column, second row – blue area). For A20 feedback strengths above 1, an increase in the A20 feedback strengths can lead to a delay in the time of the maximal concentration of NF-κB. Very high A20 feedback strengths completely diminish the NF-κB response. The effect of the A20 feedback on the response time of NF-κB is also modulated by the IκBα feedback (Fig 4C – first column, third row). The increase in the response time of NF-κB for confined combinations of low A20 and IκBα feedback strengths is due to a prolonged higher concentration of NF-κB at later time points. The response time of NF-κB remains low for a wide range of different A20 feedback strengths for IκBα feedback strengths above 1. To summarize, the effects of the two feedbacks, A20 and IκBα, in model A can be subdivided into three main areas. The first area comprises combinations of A20 and IκBα feedback strengths below 1. Those combinations result in a rapid but prolonged first peak of NF-κB and a higher NF-κB concentration at later time points similar to case II in Fig 4B. The second area is determined by high A20 feedback strengths, where the NF-κB response is completely inhibited for low IκBα feedback strengths similar to case I in Fig 4B. However, if the IκBα feedback strength is high, NF-κB remains responsive. The third area comprises high IκBα feedback strengths resulting in a slightly decreased first peak of NF-κB and no response at later time points similar to case III and IV in Fig 4B.

In model B, the A20 feedback strength hardly influences the height and time of the maximal concentration of NF-κB. Both measures are mainly determined by the IκBα feedback strength (Fig 4C – second column, first and second row). However, the A20 feedback strength influences the response time of NF-κB (Fig 4C – second column, third row). Especially, if the A20 and IκBα feedback strengths are both low, the NF-κB response time is higher. Thus, in model B the initial NF-κB response is mainly determined by the IκBα feedback, whereas the combination of both feedbacks influences the NF-κB dynamics at later time points.

In model C, an increase in the A20 feedback strength reduces the maximal concentration of NF-κB for A20 feedback strengths above 1 (Fig 4C – third column, first row). For feedback strengths below 1, the A20 feedback barely influences the maximal concentration of NF-κB. In those cases, an increase in the IκBα feedback strength can gradually decrease the maximal concentration of NF-κB. The time of the maximal concentration of NF-κB appears to be mainly robust towards changes in the two feedback strengths (Fig 4C – third column, second row). Only combinations of A20 feedback strengths above 1 and IκBα feedback strengths below 0.1 delay the time of the maximal concentration of NF-κB. Considering the response time of NF-κB, the influence of the A20 feedback can be strongly modulated by the IκBα feedback (Fig 4C – third column, third row). The NF-κB response time remains low for IκBα feedback strengths above 1 independent of the A20 feedback strength. For an IκBα feedback strength below 1, the A20 feedback strength can increase the NF-κB response time for A20 feedback strengths either above 10 or for feedback strengths between 1 and 0.1. To summarize, the effects of the two feedbacks in model C can be subdivided into three areas. The first area comprises combinations of A20 and IκBα feedback strengths below 1. Those combinations result in a rapid, but prolonged first peak of NF-κB and a higher NF-κB concentration at later time points similar to case II in Fig 4B. The second area is confined by A20 feedback strengths above 10 and IκBα feedback strengths below 0.1 resulting in a reduced as well as a delayed maximal NF-κB concentration similar to case I in Fig 4B. The third area comprises IκBα feedback strengths above 1 leading to a fast but decreased first peak of maximal NF-κB and no response at later time points similar to case III and IV in Fig 4B.

Altogether, the models show similar, but also different influences of the feedbacks on the NF-κB dynamics. For model A and C, the two negative feedbacks, IκBα and A20, have an impact on the initial dynamics. Both can independently reduce the maximal NF-κB concentration. However, in both models the two feedbacks are not completely redundant but have distinct functions in modulating the NF-κB response. If both feedback strengths are below 1, the inhibitory effect of A20 and IκBα is weak. In that case, the initial NF-κB response is slightly delayed and a prolonged activation of NF-κB can be observed at later time points. If A20 feedback strengths are high, the NF-κB response is completely inhibited in model A. In model C, a reduced as well as delayed NF-κB response can be observed. If the IκBα feedback strength is high, both models show a reduced but fast initial NF-κB increase and no response at later time points. To summarize, in models A and C both feedbacks inhibit the maximal concentration of NF-κB, but the A20 feedback delays the initial response and prolongs the response at later time points, whereas the IκBα feedback results in a faster initial activation and rapid deactivation of NF-κB. In contrast, in model B the initial NF-κB response is hardly influenced by the A20 feedback but mainly regulated by the IκBα feedback. Also in model B both feedbacks have an effect on the later phase of the NF-κB dynamics.

### Characterization of the interplay of TNFα stimulation and A20 feedback strengths

In all three considered mechanisms, the A20 feedback modulates the signal transduction of the TNFα stimulus towards the activation of IKK. We are therefore interested in the influence of the A20 feedback strength on the NF-κB response upon different strengths of TNFα stimulation. To address this question, we simultaneously varied the stimulation strength of TNFα and the strength of the A20 feedback and quantified their influence on the maximal concentration of NF-κB, time of the maximal concentration and the response time of NF-κB (Fig 5). Here, the IκBα feedback strength is fixed to the value of 1.

**Fig 5:**
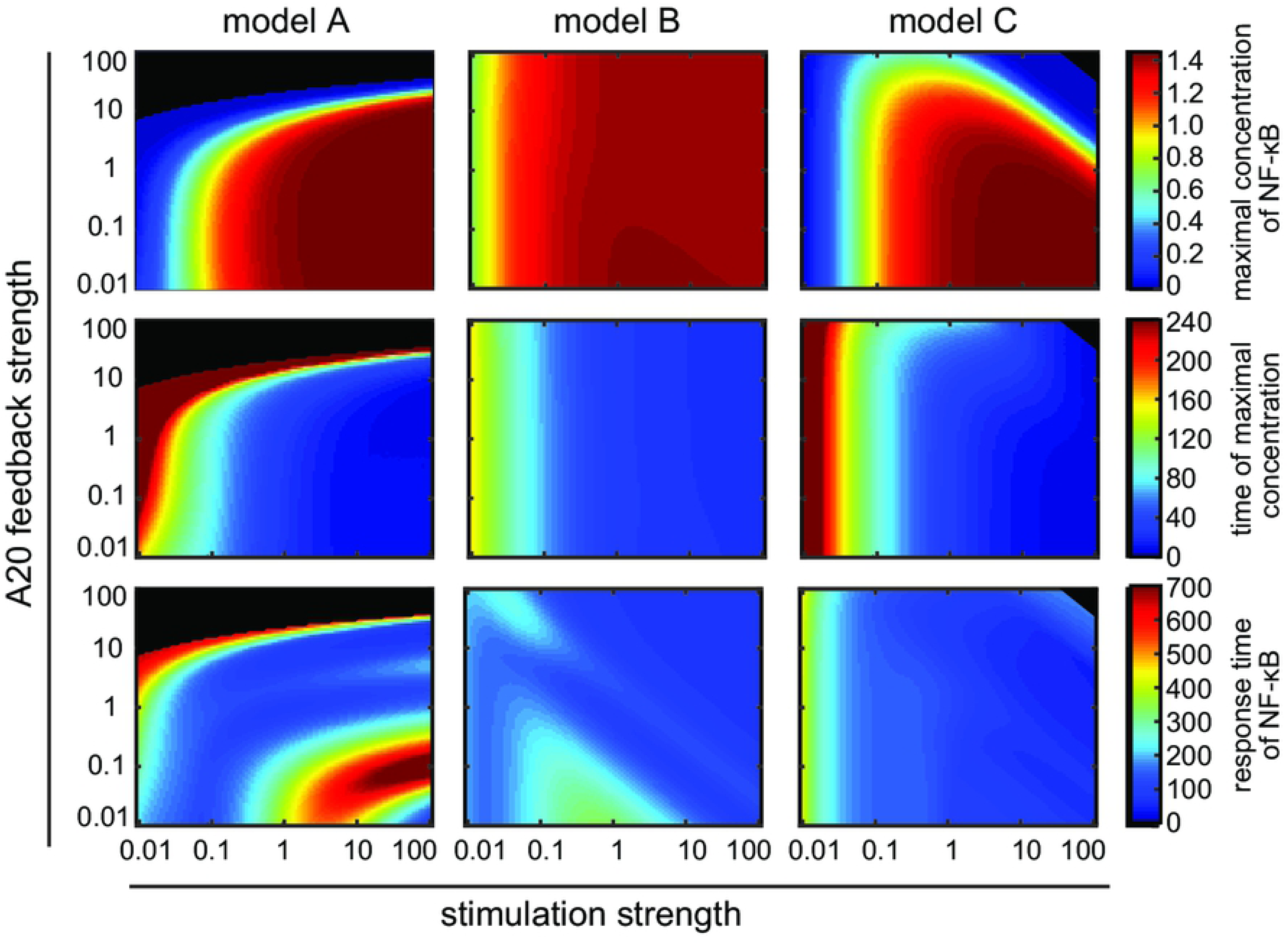
Influence of A20 feedback strength and TNFα stimulation strength on NF-κB dynamics. NF-κB dynamics of model A (first column), model B (second column) and model C (third column) are characterized by the maximal concentration of NF-κB (first row), the time of the maximal concentration of NF-κB (second row) and the response time of NF-κB (third row). Black areas mark combinations of A20 feedback strength and TNFα stimulation strength with hardly any observable NF-κB response; the difference between maximal and initial NF-κB concentrations is less than 0.001 μM.

In model A, variations in TNFα stimulation change the initial and long term dynamics of NF-κB (Fig 5 – first column). In particular, an increase in TNFα stimulation strength leads to a faster and stronger increase in the maximal NF-κB value (Fig 5 – first column, first and second row). This effect can be strongly modulated by the A20 feedback: for feedback strengths above 1 a reduction and delay of the maximal NF-κB concentration can be observed. High A20 feedback strengths above 10 result in a complete prevention of the NF-κB response for various TNFα stimulation strengths (Fig 5 – first column, black area). The response time of NF-κB is influenced by TNFα stimulation and A20 feedback strengths in a complex way (Fig 5 – first column, third row). For instance, for the combination of A20 feedback strengths below 1 and TNFα stimulation strengths above 1 the response time of NF-κB increases, indicating a prolonged NF-κB activation. In contrast, the combination of A20 feedback strengths around 0.01 and TNFα stimulation strengths above 10 leads to a decrease in the response time of NF-κB. The underlying reason is the change in the deactivation of NF-κB. For A20 feedback strengths of 0.01 and TNFα stimulation strengths of 100, NF-κB is not deactivated. Thus, NF-κB concentration does not decrease after its initial increase, resulting in a low response time (Fig 9 in S1 File). However, for A20 feedback strength of 0.1 and TNFα stimulation strengths of 100, NF-κB concentration slowly decreases after its initial increase, resulting in a high response time (Fig 9 in S1 File).

In model B, the amount and time of the maximal concentration of NF-κB depend on the TNFα stimulation strength, but are mostly robust toward changes in A20 feedback strength (Fig 5 – second column, first and second row). However, both TNFα stimulation strength and A20 feedback strength affect the response time of NF-κB (Fig 5 – second column, third row). The effect is non-linear: low TNFα stimulation strengths between 0.1 and 1 and very low A20 feedback strengths below 0.1 show an increase in the response time of NF-κB, indicating a prolonged activation of NF-κB. However, in the case of TNFα stimulation strengths between 10 and 100, a decrease in the response time is observed.

In model C, the maximal concentration of NF-κB and the timing of its peak mostly depend on TNFα stimulation strengths (Fig 5 – third column, first and second row). A20 feedback strength can lead to a reduction and a slight delay of the maximal NF-κB concentration for high TNFα stimulation strengths. In particular, if A20 feedback strength as well as TNFα stimulation strength are high, the maximal concentration of NF-κB decreases and can result in a complete prevention of the NF-κB response (Fig 5 – third column, black area). The response time of NF-κB mainly depends on TNFα stimulation strength and hardly on A20 feedback strength (Fig 5 – third column, third row).

In conclusion, the initial dynamics, that is the maximal NF-κB concentration and its timing, are strongly determined by the TNFα stimulation strength in all models. In models A and C the A20 feedback can strongly modify that impact. However, in model B, we see no significant effect of the A20 feedback on the amount and time of maximal NF-κB. The effect of the TNFα stimulation strength and the A20 feedback on the long-term dynamics is more complex. However, if we consider the effect of TNFα stimulation (for factors >1) and a given A20 feedback strength (factor = 1), we observe opposite effects in the models: while a higher TNFα stimulation strength leads to an increase of the response time in model A, that is the long term dynamics, such a stimulus increase would cause a decrease in the response time in models B and C.

### Comparison of simulations with experimental data for the effect of varied TNFα stimulation strength

The qualitative differences between the models suggest an experimental setup to scrutinize the A20 feedback implementations. To predict the outcome of such an experiment, we simulated the NF-κB dynamics of the models A-C in response to three different TNFα concentrations (Fig 6A). We selected TNFα stimulation because changes in TNFα concentration are easier to perform experimentally than changes in A20 feedback strength. Our simulations predict for model A that NF-κB levels remain high for stimulation with 100 ng/ml TNFα compared with 10 ng/ml TNFα at later time points (Fig 6A). In contrast, in models B and C, NF-κB levels decrease faster at later time points upon stimulation with 100 ng/ml TNFα compared to 10 ng/ml TNFα. These predictions are independent of the assumed A20 feedback strengths (Fig 10 in S1 File) and are furthermore verified by simulations of the models published by Murakawa et al. (2015) (13), Ashall et al. (2009)(21) and Lipniacki et al. (2004) (10) (Fig 11 in S1 File).

**Fig 6:**
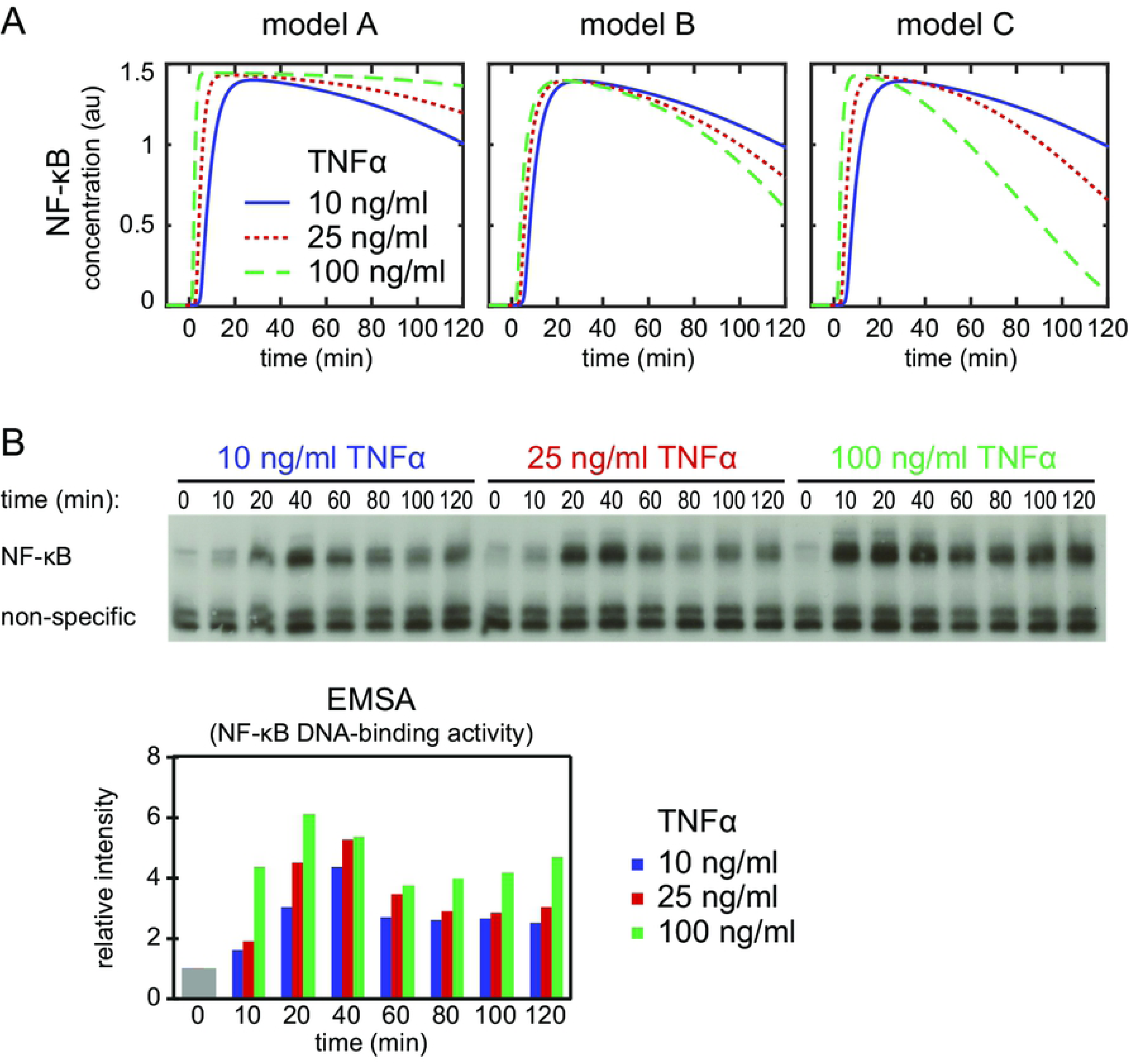
Dynamics of NF-κB upon stimulation with different TNFα concentrations. A: Simulation of NF-κB assuming a stimulation with 10 ng/ml (solid line), 25 ng/ml (dotted line) and 100 ng/ml TNFα (dashed line) in model A (left), model B (middle) and model C (right). B: Exemplary EMSA experiment measuring NF-κB DNA-binding activity over a time course of 120 min in HeLa cells upon stimulation with 10 ng/ml, 25 ng/ml and 100 ng/ml TNFα. The histogram shows the quantification of the EMSA experiment. The mean value of the relative intensities at t=0 is set to 1 and used as a normalisation for all other values. Two replicate experiments are shown as supplemental Fig 12 in S1 File.

We validated our model predictions by applying 10 ng/ml, 25 ng/ml and 100 ng/ml TNFα to HeLa cells. The time course measurements of NF-κB’s DNA-binding activity by EMSA showed NF-κB dynamics as predicted for model A but not model B or C (Fig 6B). The experiments thus indicate that the implementation of the A20 feedback structure of model A is appropriate to describe the effect of A20 on the dynamics of NF-κB in HeLa cells.

## Discussion

In this study, we developed a modular modeling approach to analyze the impact of different A20 inhibition mechanisms on the dynamics of NF-κB. In particular, we compared three distinct implementations of the A20 feedback by combining upstream modules of available models with a common core pathway module. By fitting the resulting models to a comprehensive experimental data set, we derive models with quantitatively comparable NF-κB dynamics. When analyzing the effect of variations of the strength of the A20 and IκBα feedbacks, as well as of TNFα stimulation in these models, we observe similarities, but also model-specific differences. Increasing IκBα feedback strengths attenuate the initial as well as the long-term NF-κB response in all three models, that is, reduce the maximum and response time, respectively. Increasing A20 feedback strengths reduce the maximum and duration of the NF-κB response in models A and C. In model A, the NF-κB response is even completely diminished for very high A20 feedback strengths. However, in model B the A20 feedback has no impact on the initial dynamics. Moreover, our simulations predicted that changes in the TNFα stimulation strength influence initial and long-term dynamics of NF-κB. Here, we observed qualitative differences in the long-term NF-κB response between the different models. We used these predictions for an experimental validation in HeLa cells. The experimental observations strongly support model A, but not model B or C.

Models A-C differ in the implementation of the A20 feedback. In all three models, A20 acts conjointly with the stimulus in order to inhibit IKK activation. Model A includes in addition a basal IKK activation rate that is inhibited by A20 (reaction 14). Such a composite, non-linear description of the inhibitory influence of A20 seems necessary to reproduce the NF-κB dynamics of HeLa cells. This indicates that the regulation of IKK activity by A20 in this cell type may result from a combination of several mechanisms and is thus more complex than anticipated. Indeed, A20 seems to fulfil multiple functions *in vivo*, such as a deubiquitinating activity mediated by its N-terminal ovarian tumor (OTU) domain and an E3 ubiquitin ligase activity mediated by its C-terminal zinc finger domain (5). These distinct functions of A20 may regulate the activity of upstream signal mediators and constitute potential mechanisms that may explain the complex non-linearity in the signal transduction from TNFα stimulation to IKK activation (26). A recent analysis of temperature effects on the NF-κB pathway also highlights the importance of the A20 feedback and the necessity to extend and modify its implementation in model B (21, 27). Moreover, it will be interesting to explore the role of additional negative regulators on the pathway, e.g. the deubiquitinating enzymes CYLD and OTULIN (5) as well as the effect of the cross-talk with the non-canonical pathway (21, 28, 29).

Our analyses of the three models revealed redundant but also distinct functions of the two negative feedbacks, A20 and IκBα. This confirms and extends earlier findings by Werner et al, 2008 (20), demonstrating distinct roles of the two feedbacks in a very detailed pathway model. In that publication, IκBα has been reported to modulate mostly the initial NF-κB response while A20 mainly shapes the late response. In our current study, we characterize the output based on quantitative measures for a wide range of different feedback strengths. We find that the IκBα feedback fine-tunes the initial NF-κB response in all models. However, it can also influence the response-time and therefore the long-term dynamics. The A20 feedback has different effects in models A, B and C. In models A and C, it modulates the initial as well as long-term dynamics. Moreover, in model A it has a bimodal on-off effect on the NF-κB response, i.e. preventing the NF-κB response at high A20 feedback strengths. The non-redundant functions of the two negative feedbacks could be due to their structural properties: the two feedbacks are interlocked, with the IκBα feedback serving as an inner feedback loop and the A20 feedback as an outer feedback loop. Previous studies indicted distinct functions of interlocked feedback loops with respect to the oscillatory behavior of a system (30, 31). Here, a weak or strong outer feedback loop may cause an on or off response, respectively, independent of the strength of the inner feedback loop. However, the inner feedback loop can fine-tune the response in the case of a weak outer feedback loop. Such interlocked feedback loops are very common regulatory motifs in signaling pathways in general (32-35).

Taken together, our quantitative modular modeling approach employs the regulation of NF-κB signaling by the A20 feedback as an example case to study the impact of different implementations of an inhibition mechanism on the model’s response to perturbations. Comparing the simulations of the three models A-C to experimental data suggests that model A is an appropriate choice to describe TNFα stimulation in HeLa cells. Our results emphasize the need to further explore the molecular details of processes upstream of IKK regulation.

## Acknowledgments

None.

## Supporting information

**S1 File. Supplemental material.**

Supplemental figures and tables, detailed description of the mathematical models, details on parameter estimation, sensitivity analysis, and model predictions.

